# An immunocompetent Merkel cell carcinoma model for preclinical studies

**DOI:** 10.64898/2026.06.25.734228

**Authors:** Monique E. Verhaegen, Simar Bhatia, Kathrine L. Singer, Miranda Baumbick, Pei-Wei Huang, Li-Jyun Syu, Dawn Wilbert, Avery Selig, Giselle A. Farjo, Emily Walter, Natalia Wolinski, Allison Furgal, Denise A. Galloway, Paul W. Harms, Marcin Cieslik, Andrzej A. Dlugosz

## Abstract

Merkel cell carcinoma (MCC) is a rare and aggressive neuroendocrine skin cancer that frequently carries integrated Merkel cell polyomavirus DNA and expresses oncogenic viral small T antigen (sTAg) and truncated large T antigen (tLTAg). We previously reported a mouse model of MCC with skin-targeted expression of sTAg, tLTAg, and the Merkel cell transcription factor ATOH1, combined with deletion of *Trp53*. Here, we optimized this model to achieve 100% tumor penetrance with lymph node metastases, established four mouse MCC cell lines, and selected one line, mMCC2, for pilot preclinical trials. In immunocompetent C57BL/6J mice, mMCC2 cells reliably produce MCCs and lymph node metastases following subcutaneous or intradermal (orthotopic) injection, and liver and lung metastases after tail vein injection. Mouse MCC allografts resemble parental tumors histologically and express a full complement of MCC differentiation markers. Treatment of allografted mice with anti-PD-1 resulted in variable inhibition of tumor growth. In contrast, treatment with lysine-specific histone Wdemethylase 1 (LSD1) inhibitors, with or without anti-PD-1, led to consistently lower tumor volumes by 5.7-fold in both groups (P < 0.0001) and smaller or undetectable lymph node metastases. Growth-inhibited tumors in all groups showed a marked reduction in proliferating tumor cells and increased infiltration by F4/80+ macrophages and CD8+ T cells. These findings support a role for immune-cell recruitment in treatment response and underscore the importance of immunocompetent preclinical models, even in studies using targeted therapies. This unique virus-positive MCC allograft model, which produces local tumors as well as regional and distant metastases in immunocompetent hosts, provides a critical platform for preclinical evaluation of new therapeutic strategies and sets the stage for much-needed translational studies to inform future clinical trials.

## INTRODUCTION

Merkel cell carcinoma (MCC) is a rare and highly aggressive neuroendocrine skin cancer characterized by early metastasis, high recurrence rates, and significant mortality (reviewed in (1–3)). Approximately eighty percent of MCCs are virus-positive (VP-MCC), arising through clonal integration of Merkel cell polyomavirus (MCPyV) DNA and expression of the viral oncoproteins truncated Large T antigen (tLTAg) and small T antigen (sTAg), which contribute to tumorigenesis (4). In contrast, virus negative (VN)-MCC is driven by chronic UV exposure and characterized by a high tumor mutational burden (TMB) and UV-signature mutations affecting genes encoding several oncogenic drivers (5–7).

Depending on MCC tumor stage, optimal treatment may include surgery, lymph node dissection, radiation therapy, and systemic therapy (reviewed in (8)). While surgical excision remains the standard treatment for early-stage primary tumors, immune checkpoint inhibitors (ICIs) are now first-line therapy in locally advanced, recurrent, or metastatic MCC (9–11). ICI has significantly improved treatment outcomes, with approximately 50% of patients achieving an objective response. However, many patients fail to respond to treatment, experience significant side effects, or develop refractory disease (reviewed in (8, 12–16), underscoring the need for more effective approaches to therapy.

*In vitro* studies using cell lines can identify potential treatments that directly affect tumor cells, however, *in vivo* approaches are essential for modeling cancer-host interactions, investigating various aspects of tumor biology, and testing new treatment approaches. Most *in vivo* studies on MCC have used human MCC cells or patient-derived tumor samples grown as xenografts in immunodeficient mice (reviewed in (17–19)). However, given the importance of immune cells in treatment response to ICI as well as many other forms of therapy (reviewed in (20, 21)), preclinical studies using MCCs in immunocompetent mice are an especially high priority and unmet need. The lack of a tractable, immunocompetent MCC mouse model has greatly hindered progress in translational studies needed to guide future clinical trials.

We recently developed the first virus-positive GEMM that spontaneously develops MCCs following expression of MCPyV tLTAg, sTAg, the transcription factor ATOH1, and p53 deletion (SLAP mouse) (22). However, given the requirement for an extensive breeding scheme needed to combine multiple alleles and a complex transgene induction regimen, the SLAP model is not well suited for preclinical studies. Here, we established MCC cell lines from modified SLAP mice: these cell lines consistently produce skin tumors and metastases when injected into C57BL/6J mice. We use this novel allograft model to assess MCC treatment responses to ICI, targeted therapy, or combination therapy, paving the way for much-needed translational studies for this highly aggressive tumor.

## RESULTS

### Modification and optimization of the SLAP MCC mouse model

We initially generated and validated a *K5-CreERT2;R26-LSL-rtTA;tetO-sT/tLT;tetO-Atohl;Trp53^+/fl^* GEMM. This model was treated with tamoxifen to activate Cre function, induce rtTA expression and delete one copy of *Trp53;* and with doxycycline to induce expression of MCPyV sTAg and tLTAg, and the Merkel cell lineage driver ATOH1 (22). These mice, designated SLAP for expression of **<u>s</u>**TAg, t**<u>L</u>**TAg, and **<u>A</u>**TOH1, and heterozygous deletion of **<u>p</u>**53, developed cutaneous tumors similar to human MCCs as revealed by extensive cross-species credentialing. However, our transgene induction protocol in adult mice resulted in approximately 40% of SLAP mice requiring euthanasia for humane reasons after becoming moribund or developing large internal tumors. Also notable was loss of the wild-type *Trp53* allele in all tumors that were analyzed. Given the apparent requirement for complete loss of p53 for MCC tumorigenesis we modified the SLAP model to include two floxed *Trp53* alleles (SLAP2 mouse) (**Suppl. Fig. 1a**). In addition, since low tumor penetrance may have resulted from immune elimination of tumor cells following induction of immunogenic viral antigens in adult mice, we modified our transgene induction protocol. We induced transgene expression at postnatal day 4-5 by applying topical 4-hydroxy-tamoxifen (4OHT) directly to dorsal skin of neonatal pups and providing doxycycline chow to lactating females, reasoning that induction of viral antigens at this early developmental stage would promote immune tolerance leading to higher tumor penetrance. Pups were continued on doxycycline after weaning and were monitored until tumors developed (**Fig. 1a, Suppl. Fig. 1b)**.

**Figure 1.**
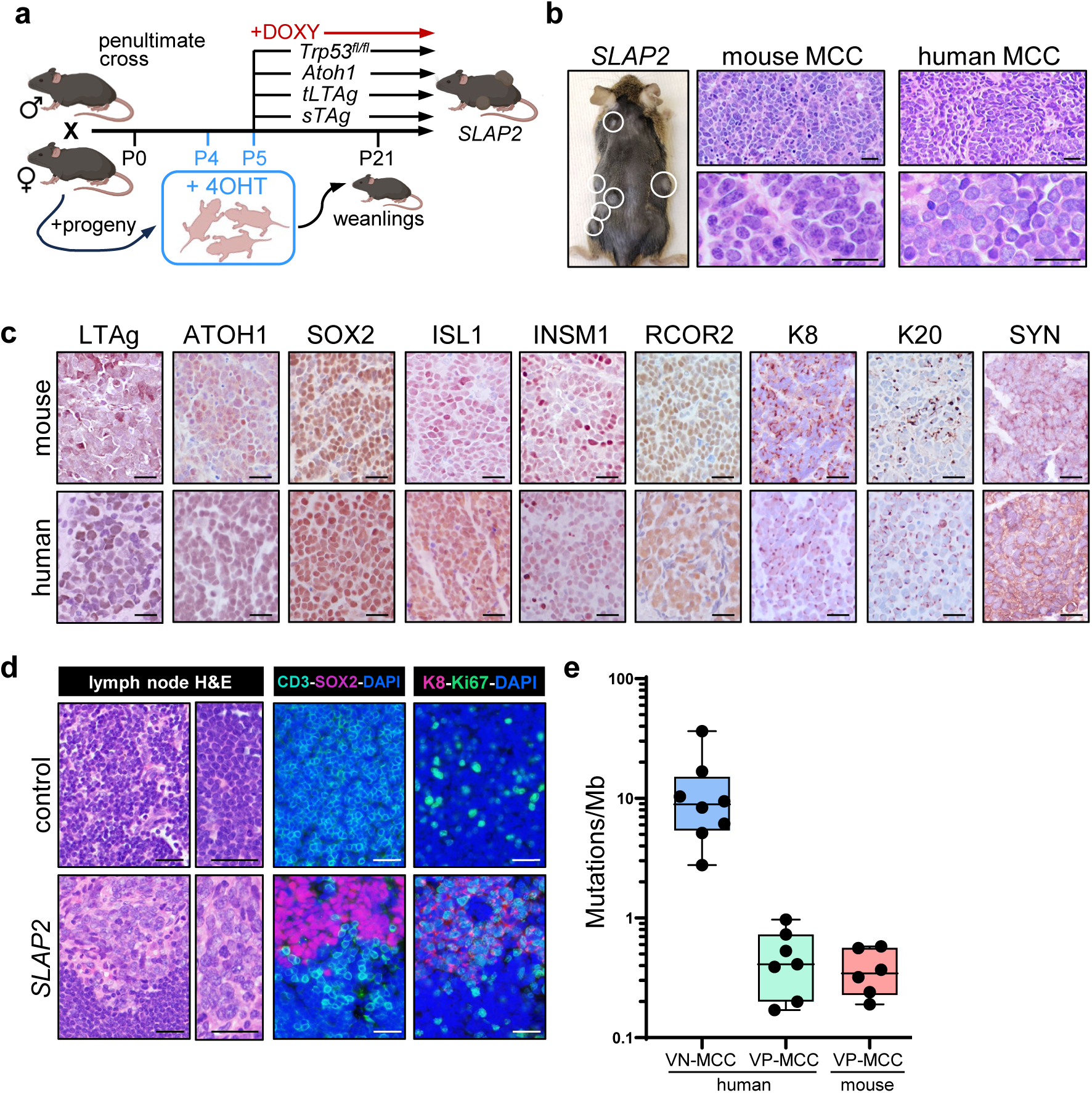
*K5-SLAP2* mice develop mouse MCC tumors and regional metastases. a) Modified breeding and transgene induction protocol (additional details in text). b) Gross image of tumors on dorsal skin of SLAP2 mouse. H&E images show similar morphology of mouse and human tumors. c) IHC for MCPyV LTAg and MCC markers in mouse and human tumors. d) Regional lymph nodes in SLAP2 mice contain highly proliferative Ki67+/SOX2+/K8+ tumor cells. The pan-T-cell marker CD3 specifies T lymphocytes in the LN. e) Tumor mutational burden is low in both mouse and human VP-MCC, with markedly higher levels in human VN-MCC. All scale bars = 25µm.

### SLAP2 mice develop dorsal skin MCC tumors and regional lymph node metastases

Biopsies of mice 3 weeks post transgene induction showed multiple hair follicle-associated tumors (**Fig. 3a**) that express all MCC markers examined (data not shown). While most microscopic tumors disappeared by 2 months, some developed into macroscopic cutaneous tumors (**Fig. 1b**). This refined modeling strategy yielded 100% tumor penetrance in all SLAP2 mice (n=11; 6F/5M), which developed macroscopic skin tumors between 8 and 16 wks of age. Tumor-bearing mice appeared bright and healthy with mice developing 1-7 skin tumors at the time of harvest, and approximately 20% also developing internal tumors. Additional skin tumors might have developed if mice were monitored for longer periods. Two mice were sacrificed early due to actively bleeding tumors from possible ulceration or trauma.

Tumors arising in SLAP2 mice closely resembled human MCCs (**Fig. 1b**), exhibiting a monomorphic small blue cell morphology with finely stippled chromatin, scant cytoplasm, and nuclear molding, with less morphological heterogeneity than the original SLAP tumors (22). All SLAP2 skin tumors were deemed histologically consistent with human MCC by a board-certified dermatopathologist (PWH). Immunohistochemical (IHC) staining for transgenes (LTAg, ATOH1) and MCC markers including SOX2, ISL1, INSMI, RCOR2, Keratin (K) 8, K20 and Synaptophysin (SYN) were comparable to human virus-positive tumors (**Fig. 1c**). Interestingly, the late-stage markers K20 and SYN were not present in MCCs arising in the original SLAP model.

Furthermore, microscopic metastases were detected in regional lymph nodes of tumor-bearing *K5-SLAP2* mice at the time of harvest. Although LNs did not always appear visibly enlarged, immunostaining revealed foci of proliferative Ki67+/SOX2+ MCC tumor cells that expressed K8 in a pathognomonic dot-like pattern (**Fig. 1d**).

### SLAP2 MCCs and human VP-MCCs have low tumor mutational burden

Whole exome sequencing was performed to analyze the tumor mutational burden (TMB) of mouse MCC tumors compared to human MCCs. TMB was defined as the number of somatic mutations per megabase of coding DNA analyzed and reported as Mutations/Mb. Tumors arising in SLAP2 mice exhibited low TMB, in the same range as human VP-MCCs; in contrast, human VN-MCCs had a markedly higher TMB (**Fig. 1e**), in keeping with prior studies (5–7).

Additionally, the fraction of genome altered (FGA), which is a metric of copy-number alteration, was in the range of 3.5-12.1% (data not shown), indicating a relatively stable genome.

### Establishment of mouse MCC cell lines

Mouse MCC cell lines were established in a manner similar to our previously described human UM-MCC cell lines (23). Mouse lines grew in suspension as aggregates and were supplemented with fresh medium or passaged every 2-3 days once established. In total, we established four mMCC lines, from independent SLAP2 tumors that represented classical MCC-like histology and expressed all MCC markers, that went on to be tested for tumorigenicity in C57BL/6J mice. Representative cell lines designated mMCC1 and mMCC2, established from tumors arising in two different SLAP2 mice, resemble human MCC cell lines *in vitro* (**Fig. 2a**). Immunoblots verify expression of transgene-derived LTAg and ATOH1 in cell lines mMCC1 and mMCC2, with comparable levels of LTAg expressed in the human VP-MCC cell line UM-MCC52 (**Fig. 2b**).

**Figure 2.**
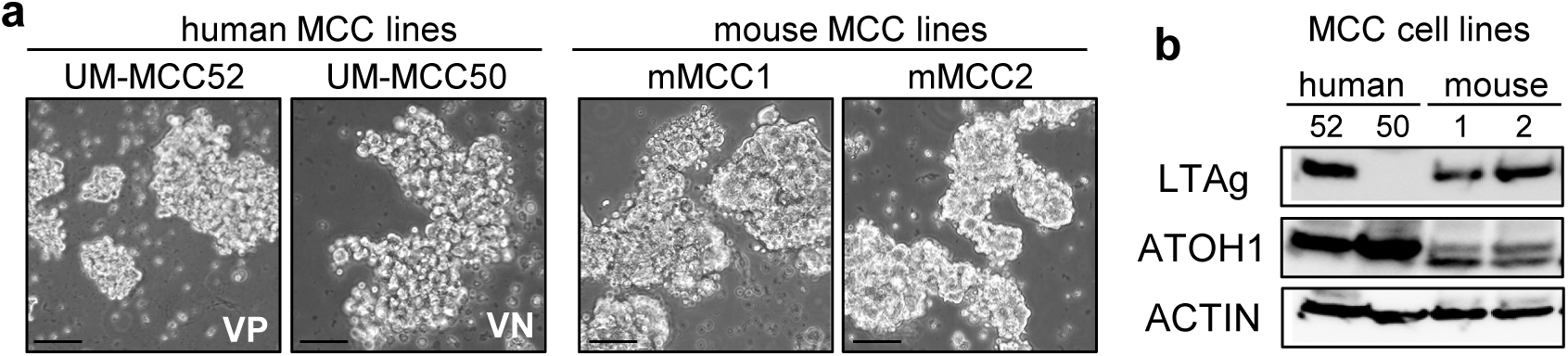
Representative MCC cell lines. a) Phase-contrast images showing virus positive (VP) and virus negative (VN) human MCC cells and representative VP mouse MCC cells derived from SLAP2 tumors, each growing as aggregates in suspension. Scale bars = 50µm. b) Immunoblotting for MCPyV LTAg, ATOH1, and actin in lysates of human and mouse MCC cell lines shown in a).

### Tumorigenicity and lymph node metastasis of mouse MCC cell line allografts in immuncompetent mice

To develop an allograft mouse model that would be useful for preclinical studies requiring an intact immune system, we tested four mMCC cell lines for tumorigenicity in C57BL/6J mice. Following subcutaneous (SQ) injection, two lines (mMCC1 and mMCC2) grew with 100% penetrance, one with ∼50% penetrance, and one failed to grow. This variable penetrance may reflect partial immune incompatibility, since the primary tumors used to establish these cell lines arose in SLAP2 mice that were not fully congenic on the C57BL/6J background (see Methods).

Since human MCCs typically reside in the dermis (24), as recapitulated in the SLAP mouse model (22) and shown here for early and full-blown SLAP2 mouse tumors (**Fig. 3a)**, we tested tumorigenicity of mouse cell lines in an orthotopic setting. Of the two lines that grew consistently as SQ tumors, only mMCC2 cells reliably produced tumors following intradermal (ID) injection. This difference in tumor take-rate is in keeping with prior studies showing a more permissive immune microenvironment in SQ than ID sites (25, 26), with varying degrees of immune incompatibility of different cell lines a likely contributor. Allograft tumors grew to approximately 100 mm^3^ in 2-3 weeks, and mice required euthanasia in 7-8 weeks due to tumor burden (>2cm) (see vehicle-treated mice in **Fig. 5a**).

**Figure 3.**
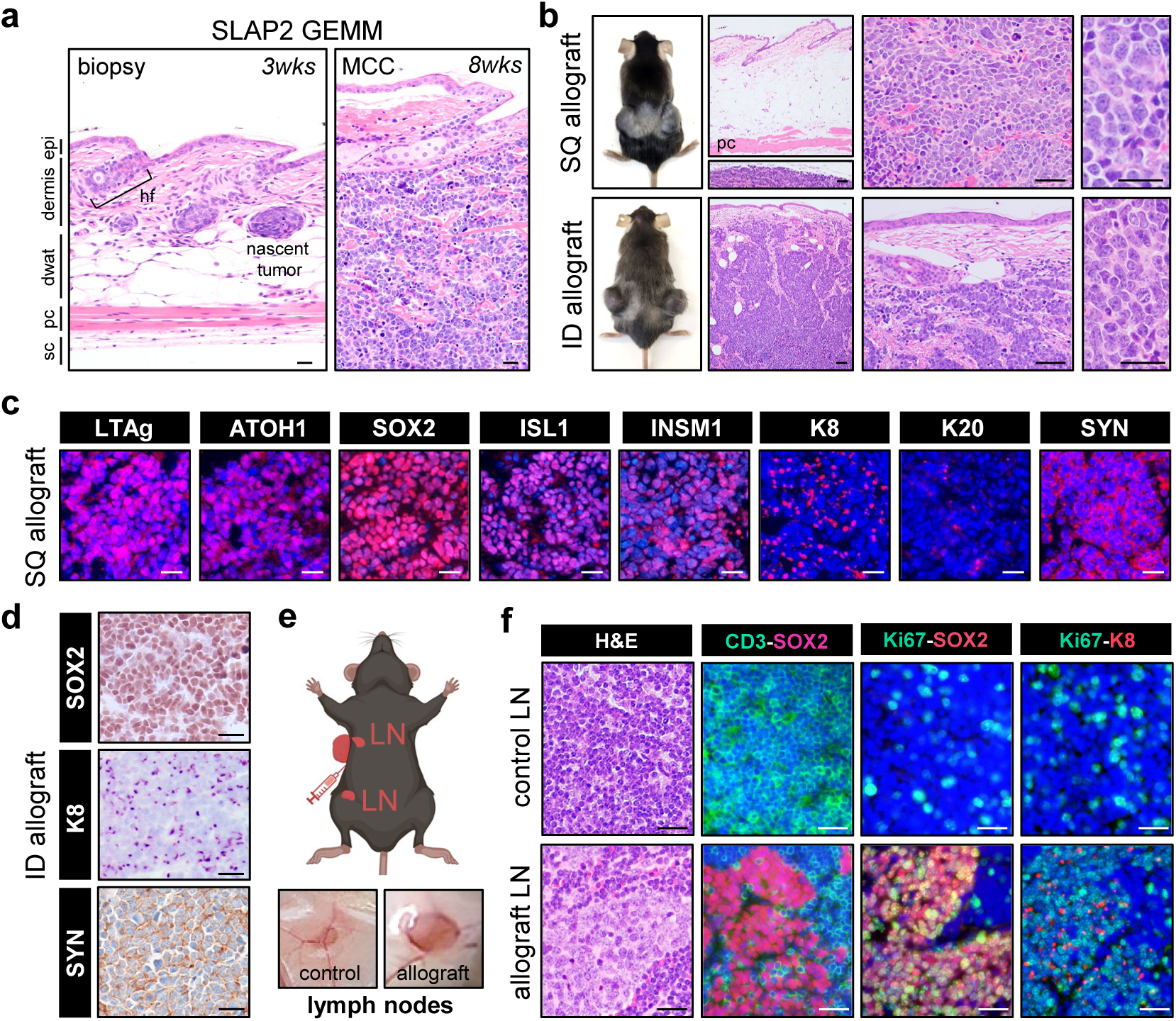
Mouse MCC cell line growth in immunocompetent mice mimics autochthonous tumors. a) Histology showing location of arising nascent tumors at 3 wks (left panel) or a full blown MCC tumor at 8 wks (right panel) post transgene induction in the dermis of a SLAP2 mouse. hf = hair follicle, epi = epidermis, dwat = dermal white adipose tissue, pc = panniculus carnosus, sc = subcutis. b) Representative whole-body images of mice with SQ or ID allografts at 5.5wks post cell implantation. Histology depicts the appearance of tumor cells below the panniculus carnosus or in the dermis, for SQ and ID tumors, respectively. The SQ allograft tumor was located far below the pc. c) Co-IF staining of OCT-embedded frozen sections of SQ allograft tumors reveal expression of transgenes (LTAg and ATOH1) and the full panel of MC/MCC lineage markers. d) Immunohistochemical staining for early- (SOX2), intermediate-(K8), and late-stage (synaptophysin/SYN) Merkel cell/MCC lineage markers in orthotopic ID tumor allografts. e) Mice develop enlarged regional LNs following ID injection of mMCC2 cells.f) Histological analysis by H&E and co-IF staining shows the presence of CD3+ T cell lymphocytes and proliferative Ki67+/SOX2+/K8+ MCC tumor cells in LNs. All scale bars = 25µm.

The mMCC2 allografts in C57BL/6J recipients (**Fig. 3b**), and the primary mouse tumor from which this cell line was derived (shown in **Fig. 1b**), exhibited strikingly similar MCC-like histology. SQ allograft tumors were located below the panniculus carnosus muscle layer while ID allografts developed within the dermis (**Fig. 3b**), where most human MCCs arise.

Immunofluorescent (IF) staining of frozen sections from SQ mMCC2 allografts revealed the expected expression of transgene-derived LTAg, ATOH1, and multiple MCC markers (**Fig. 3c**) as also seen in the parental SLAP2 tumor (**Fig. 1c**). Confirmation of early, intermediate, and late-stage MC/MCC markers was additionally confirmed in ID mMCC2 allografts by IHC (**Fig. 3d**).

Strikingly, mice with mMCC2 allograft tumors consistently developed regional lymph node metastases (**Fig. 3e**). All inguinal LNs were visibly enlarged in mice with mid-size tumors (approximately 500mm^3^; N>30) and contained large areas with MCC-like histology (**Fig. 3e-f**). Some mice also displayed enlarged axillary LNs at the time of tumor harvest (data not shown). Staining revealed that all enlarged LNs showed effacement and the presence of proliferative Ki67+/SOX2+ MCC cells with characteristic dot-like K8 expression (**Fig. 3f**).

### Mouse MCC tumor cells develop visceral metastases in immunocompetent mice

No macroscopic or microscopic metastases were detected in internal organs of mice bearing mMCC2 allografts, possibly because animals required euthanasia approximately eight weeks post injection due to heavy tumor burden, limiting the time available for distant metastasis to develop. In an effort to allow longer survival, we excised allografts soon after tumor detection, but these mice also required euthanasia within 3-4 weeks post resection due to massive enlargement of regional lymph nodes. We therefore performed a well-established experimental metastasis assay by injecting mMCC2 cells intravenously (IV) into the tail vein (**Fig. 4a**) of C57BL/6J recipients. Although this approach bypasses primary tumor establishment, local invasion, and intravasation, it enables assessment of tumor cell survival in the circulation, extravasation, and colonization of distant organs (27). Six weeks following IV injection of tumor cells mice were euthanized due to marked abdominal distension. Necropsy of injected mice revealed livers with numerous macroscopic tumor nodules and liver weights on average 4-fold higher than controls (**Fig. 4b**). While lungs of tail vein injected mice appeared similar in size to control lungs, several minute nodules were identified. Histologic evaluation revealed large foci of MCC-like tumor cells in livers and much smaller foci in lungs; in all mice analyzed (N=7), these cells were highly proliferative (Ki67+) and expressed MCC markers SOX2 and K8 in a dot-like pattern (**Fig. 4c-d**).

**Figure 4.**
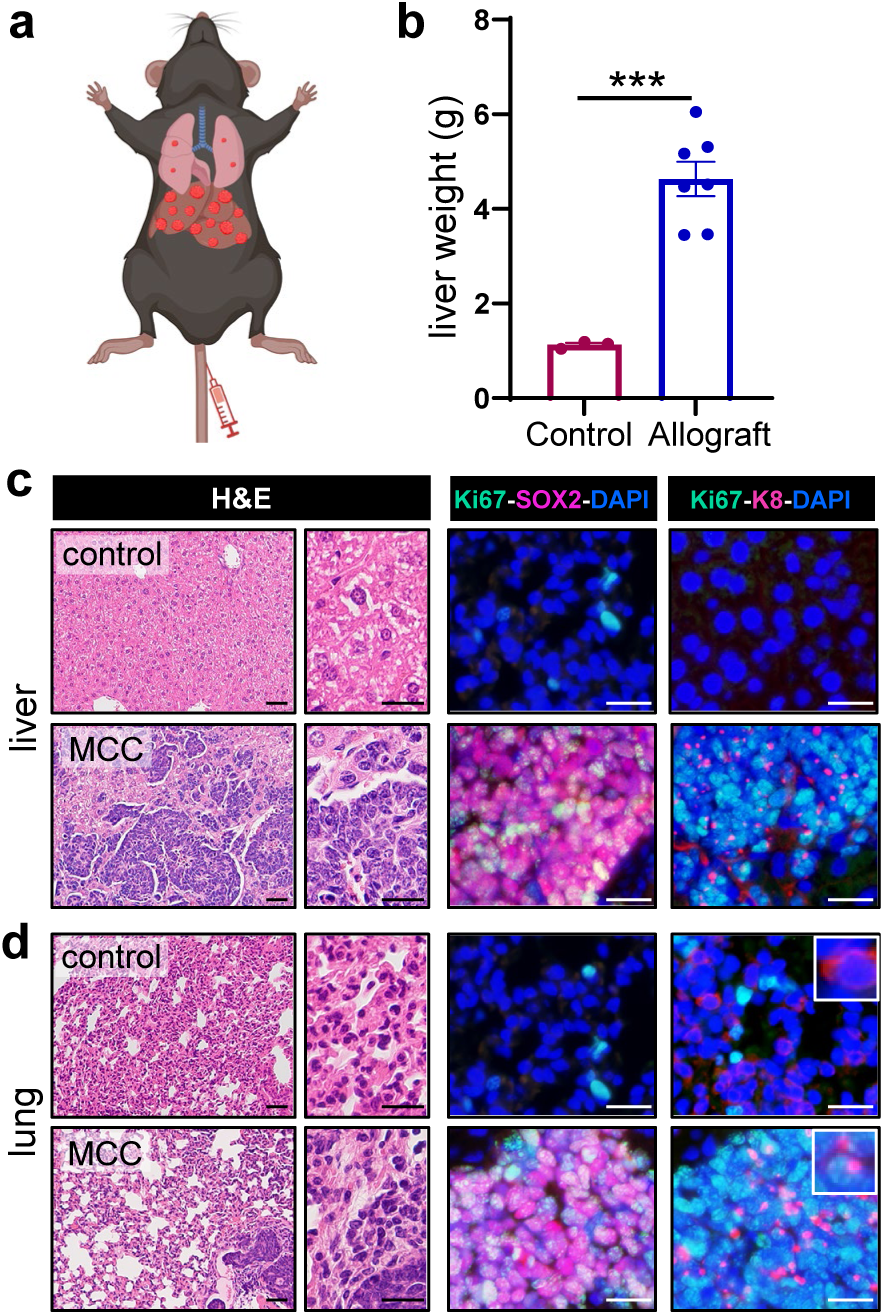
Development of experimental mouse MCC metastases in liver and lung. a) Schematic of experimental assay for distant metastases with representation of tumors arising in liver and lungs (red circles) following tail vein injection of 1x10^6^ mMCC2 cells. b) Livers contained numerous nodular tumors in cell line injected mice and were ∼4-fold greater in weight compared to controls. c) Histological analysis by H&E and IF staining confirmed multiple SOX2+ and K8+ proliferative (Ki67+) tumors in the liver. d) Histological analysis of lungs revealed smaller metastases that were highly proliferative (Ki67+) and stained for MCC markers SOX2 and K8. While normal lungs contain scattered K8+ cells, the typical dot-like pattern of K8 was detected only in MCC metastases (insets). Low mag H&E scale bars = 50µm. All other scale bars = 25µm.

### Detection of MCPyV serum antibodies in mouse models

Serum antibodies against MCPyV viral TAgs are a marker for human MCC tumor burden and can be an early predictor of tumor recurrence (28, 29). In preliminary studies, a subset of SLAP and SLAP2 mice, as well as C57BL/6J mice with mMCC2 allografts or distant metastases following tail vein injections, were seropositive at the time of euthanasia (**Suppl. Fig. 3**).

Reactivity to large and small TAgs was greatly reduced when mouse serum samples were pre-incubated with a common TAg-specific monoclonal antibody (data not shown). These results indicate that VP-MCC-bearing mice, like patients with VP-MCC, mount a humoral response to viral TAg expression.

### Response of mouse allograft MCCs to ICI, targeted therapy, and combination therapy

Given the lack of immunocompetent MCC mouse models, we next tested whether tumors arising from mMCC2 allografts would provide a useful platform for preclinical studies that include immune-modulating agents, a cornerstone of cancer therapy. Since ICI is the standard of care in locally advanced, recurrent, or metastatic MCC (9–11), we evaluated mouse MCC tumor responses to anti-PD-1 ICI. We also tested tumor responses to targeted therapy using an LSD1 inhibitor, since these drugs kill cultured human MCC tumor cells and inhibit growth of MCC xenografts (30, 31), and to anti-PD-1 combined with LSD1 inhibitor. C57BL/6J mice carrying mMCC2 allografts were treated with vehicle (N=9), anti-PD-1 (N=6), the LSD1 inhibitor bomedemstat (N=9), or anti-PD-1 plus bomedemstat (N=9). Treatment was started three weeks after ID injection, when tumors had achieved an average volume of 150 mm^3^ (**Fig. 5a**, black arrow).

**Figure 5.**
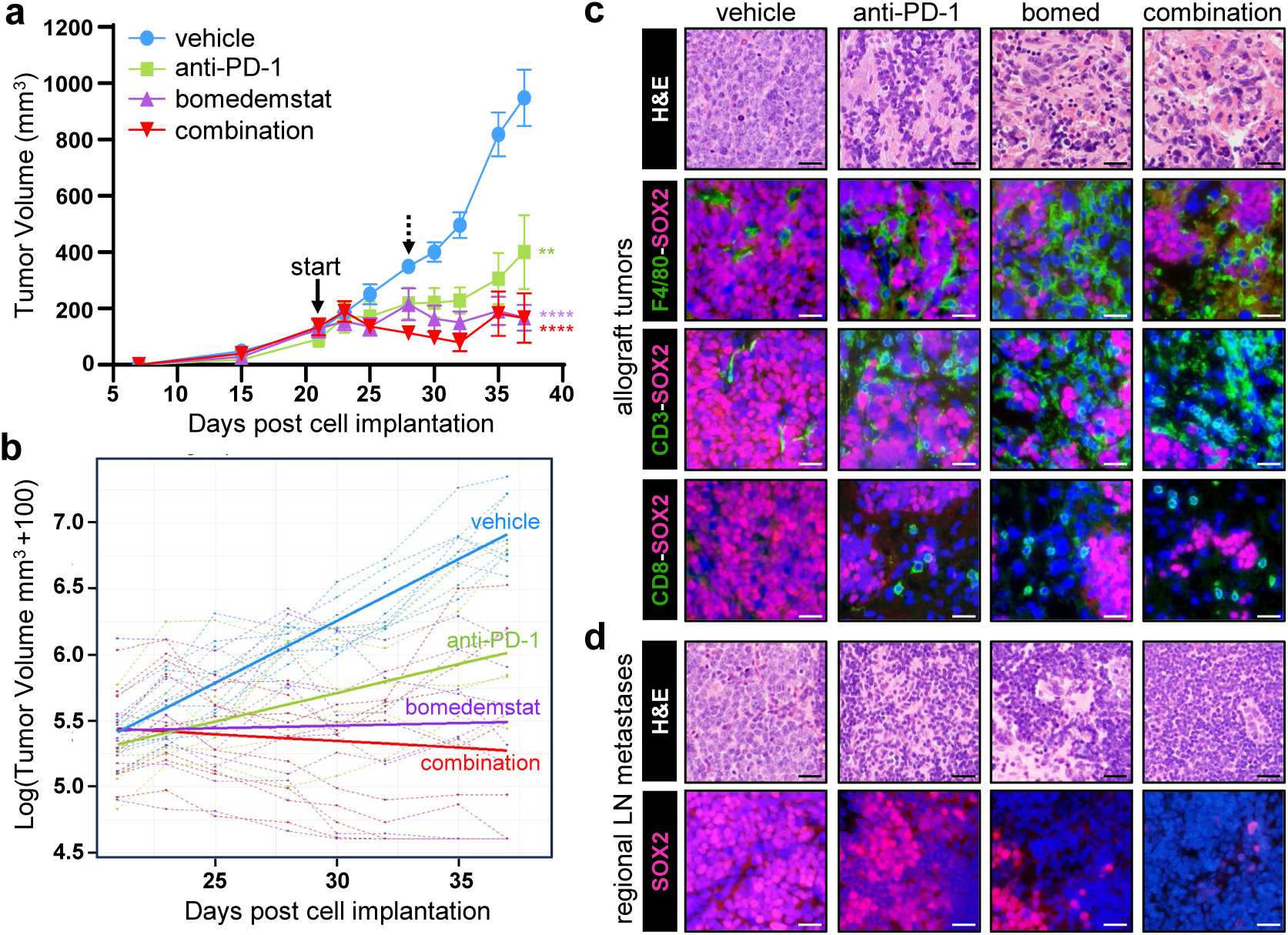
Growth inhibition of orthotopic mouse MCC allograft tumors in immunocompetent mice treated with ICI +/- targeted therapy. a) Growth kinetics of mMCC2 allografts in C57BL/6J mice following treatment with vehicle (N=9), anti-PD-1 (N=6), bomedemstat (N=9), and combination (N=9). Black arrows indicate treatment start date and one week post (dashed arrow). b) Analysis of treatment groups using a linear mixed model that estimates longitudinal growth rate (see text for additional details). c) Responders in all groups displayed decreased cellularity and a dramatic increase in immune cells including F4/80+ macrophages, CD3+ T cells, and CD8+ cytotoxic cells compared to vehicle-treated. d) Representative H&E and IF staining depict treatment response in regional LN metastases, with loss of SOX2+ MCC tumor cells in all treatment groups, particularly in combination-treated mice. Scale bars = 25µm.

Following one week of treatment, tumor volumes increased 2.8-fold in vehicle-treated mice, with an increase of 2.4-fold and 1.7-fold in anti-PD-1 and bomedemstat-treated mice, respectively. In contrast, combination treated mice displayed a 0.8-fold change in tumor volume, representing an approximate 20% decrease in tumor volume compared to starting volumes. (**Fig. 5a**, dashed arrow). Over the following 16 days of treatment, tumor growth was significantly inhibited in all drug-treated groups compared with vehicle controls. At the study endpoint, mean tumor volumes were 948.6 mm³ for vehicle, 400.1 mm³ for anti-PD-1, 167.7 mm³ for bomedemstat, and 165.8 mm³ for the combination group (**Fig. 5a**). These correspond to tumor growth inhibition of approximately 58% with anti-PD-1 (P < 0.001), 82% with bomedemstat (P< 0.0001), and 83% with combination treatment (P < 0.0001) relative to vehicle. At the study endpoint, two tumors out of nine in the bomedemstat-treated group and three tumors out of seven in the combination-treated group appeared to have completely regressed, and no visibly detectable tumor was observed. One combination-treated mouse was moribund on day 32 (N=2 tumors) and was euthanized, which may have been a result of treatment-related toxicity or repeated gavage-related tissue injury.

Regardless of the end-of-study mean tumor volumes, statistical analysis via a linear mixed model (32, 33) allowed for analysis of tumor growth rates (slope of the lines) between all groups over time. Data is shown for each individual tumor (dashed lines) and the estimated fitted line (solid line) from the linear mixed model for each group (**Fig. 5b**). Based on this longitudinal analysis, the log-volume change per day (daily rate change) was estimated to increase at 9.4%, 4.3%, and 0.39% in the vehicle, anti-PD-1 and bomedemstat-treated groups, with a 1.0% decrease per day in the combination-treated group. There were significant differences in tumor growth rates of anti-PD-1 (P = 0.0013), bomedemstat (P < 0.0001), and combination (P < 0.0001)-treated groups compared to vehicle-treated mice.

Interestingly, the degree of tumor growth inhibition in response to anti-PD-1 varied substantially across multiple experiments, reflecting the heterogeneous responses to ICI seen in patients with MCC. Although anti-PD-1 treatment yielded a statistically significant reduction in mean final tumor volume (P = 0.01, **Fig. 5a**), this result was not consistent across experiments, likely due to the variable responses among different mice (see spaghetti plots from independent experiment in **Suppl. Fig. 2**).

Histologic evaluation by H&E staining revealed decreased cellularity of tumors in all treatment group responders, most dramatically visualized in bomedemstat and combination-treated groups, as well as an apparent influx of immune cells (**Fig. 5c**). Co-IF staining revealed an increase in F4/80+ macrophages, CD3+ T cells, and CD8+ cytotoxic T cells compared to tumors from vehicle-treated mice (**Fig. 5c**). Interestingly, co-IF staining also showed a dramatic influx of immune cells following LSD1 inhibition alone. Furthermore, regional LN involvement in allografts was substantially reduced following anti-PD-1 and bomedemstat treatment, with combination therapy yielding the smallest LNs and only rare SOX2+ tumor cells detected in some LNs (**Fig. 5d**).

## DISCUSSION

Tractable mouse models that faithfully mimic human disease are essential both for basic and preclinical studies needed to advance new treatment approaches into the clinic. Despite the discovery of Merkel cell polyomavirus as a cause of human MCC in 2008 (4), the lack of immunocompetent MCC mouse models has been a major impediment to research on this aggressive malignancy. Here, building on our initial SLAP mouse model (22), we document optimization and characterization of SLAP2 mice; establishment of tumorigenic mouse mMCC cell lines; production of MCCs and regional LN metastases in immunocompetent mice carrying orthotopic mMCC cell line allografts; formation of liver and lung metastases following tail vein injection, reflecting the organotropism of human MCC metastases (34); induction of a humoral responses to viral TAgs in GEMM, allograft, and experimental metastasis models; and proof-of-concept preclinical trials using ICI, targeted therapy, and combination therapy. The availability of mMCC cell lines will provide the field with much-needed tools to advance our understanding of MCC biology, including mechanisms of metastasis, and will enable essential preclinical studies to guide future clinical trials.

Preclinical models for drug discovery have relied heavily on SQ xenografts or allografts using either human or mouse cell lines, respectively. However, the anatomical site where tumors reside influences both immunogenicity and therapeutic response, underscoring the importance of orthotopic allograft models for evaluating preclinical treatment outcomes (25, 26, 35, 36). Thus, our mouse MCC allograft model that yields both orthotopic skin tumors and regional LN metastases, as well as experimental liver and lung metastases following tail vein injection, provides the first immunocompetent MCC model capable of assessing therapeutic responses of primary as well as metastatic disease in the correct tissue context.

While immunotherapy has greatly improved outcomes in MCC patients, the durable response rate of 50% underscores the urgent need for additional MCC treatment approaches, especially for those individuals who fail ICI (37–39). In our allograft model, responses to single-agent anti-PD-1 treatment varied substantially among animals within the same treatment group. This is perhaps not surprising, since therapeutic responses to ICI in preclinical mouse models can be affected by differences in the tumor-immune microenvironment due to diet, temperature, and housing conditions (40, 41); gut microbiome composition (42–44); inter-individual heterogeneity and variable T-cell receptor repertoires, even among genetically identical mice (45, 46); and depth of tumor implantation (35).

We were particularly interested in evaluating LSD1 inhibitors in our first-in-field immunocompetent MCC model for several reasons. In prior studies, these drugs suppressed human MCC xenograft growth in immunodeficient mice and promoted tumor cell differentiation, cell-cycle arrest, and cell death (30, 31). In other tumor models, LSD1 inhibitors restore expression of MHC-I (47, 48), which is frequently down-regulated in MCC (49–51), and they enhance cytotoxic T cell activity, potentiating responses to ICI (47, 52, 53). In our mouse MCC allografts, treatment with bomedemstat alone led to a marked reduction in final tumor volumes and enhanced macrophage and T cell infiltration, highlighting the critical importance of immunocompetent mouse models for evaluating targeted therapies that modulate the tumor immune microenvironment

The results presented here validate the translational relevance of our model and underscore the critical need to develop and rigorously characterize a panel of congenic mouse MCC cell lines capable of forming syngeneic tumors after allografting in immunocompetent hosts. These cell lines would provide a powerful preclinical platform for evaluating a broad range of therapeutic strategies, including targeted therapies, ICIs and other types of immunotherapies, radiation and additional physical treatment modalities, as well as rational combination strategies, as demonstrated here for immune checkpoint blockade and LSD1 inhibition. Ultimately, mouse MCC allograft models could help identify, prioritize, and advance more effective therapeutic strategies for MCC patients who have an inadequate response to currently available treatments.

## MATERIALS AND METHODS

### GEMM Mouse Models

#### Mouse Strains

The original *K5-SLAP* mouse (SLAP) that develops spontaneous mouse MCC-like tumors has already been described (22). We used a similar strategy to generate SLAP2 mice but deleted both *Trp53* mouse alleles, since all original SLAP tumors examined were shown to have lost the second *Trp53* allele. Briefly, SLAP2 mice include a hormone inducible Cre allele (*K5-CreERT2* strain) [57]; a Cre-inducible rtTA allele (B6.Cg-*Gt(ROSA)26Sor^tm1(rtTA,EGFP)Nagy^*/J from Jackson Laboratory (Stock No. 005670) (54); and *tetO*-driven effector alleles *tetO-sT/tLT* for expression of MCPyV sT and tLT (22), and *tetO-Atoh1* (55) to coax cells into the Merkel lineage. Tissue-specific *Trp53* gene deletion was achieved with the conditional B6.129P2-*Trp53 ^tm1Brn^*/J (*Trp53 ^fl/fl^)* mutant mouse (The Jackson Laboratory, Stock No. 008462) [59]. Following multiple rounds of breeding, the final *K5-CreERT2;R26-LSL-rtTA;tetO-**<u>s</u>**T/t**<u>L</u>**T;tetO-**<u>A</u>**toh1;Tr**<u>p</u>**53^flfll^* mouse designated *K5-SLAP2* enabled tissue-specific deletion of both copies of *Trp53* following Cre-mediated recombination with 4-OHT (**Suppl. Fig. 1a**). Inducible transgenes in these mice were spatially and temporally controlled by activation of Cre function with tamoxifen leading to recombination at the ROSA locus to induce rtTA expression, which enabled expression of sTAg, tLTAg and Atoh1 following treatment with doxycycline (**Suppl. Fig. 1b**).

SLAP and SLAP2 mice were generated from parental strains including *tetO-sT/tLT* and *tetO-Atoh1* mice that still contained mixed genetic backgrounds and were not fully congenic; the other strains required for crosses were acquired and maintained on a C57BL/6J background. Progeny from crosses were genotyped using transgene-specific primers as previously described (22).

#### Transgene induction, tumor development, and tissue collection

Transgene induction was performed on neonatal mouse pups at days 4-5 to activate Cre function by applying 100µg of topical 4-hydroxy tamoxifen (4-OHT; Sigma-Aldrich) in 100% ethanol directly to the lower dorsal skin on two consecutive days. The lactating mom was simultaneously treated with doxycycline in chow (1g/kg; Bioserv Inc.) to allow for rtTA function and expression of *tetO*-regulated transgenes. At weaning, all mice were placed on doxycycline chow and maintained indefinitely; transgene negative mice were used as controls or euthanized. Mice were assessed weekly until tumors developed and tumors harvested prior to reaching maximum allowable size (2cm x 2cm). Regional lymph nodes were collected at final harvests. Dorsal skin biopsies to assess early tumor development were performed at 3 weeks post transgene induction. For all survival procedures mice were anesthetized with isoflurane gas and final euthanasia performed using carbon dioxide followed by cervical dislocation. All harvested tissue was kept in chilled Hank’s Balanced Salt Solution (HBSS) during harvest. Tissues were fixed overnight in 10% neutral buffered formalin (NBF) followed by transfer to 70% EtOH prior to tissue processing and paraffin-embedding. Tissues for frozen sections were fixed for one hour on ice in 4% paraformaldehyde, followed by overnight incubation in 30% sucrose prior to embedding in Tissue Tek optimal cutting temperature (OCT) compound. Tumor or liver samples for DNA isolation were flash frozen in liquid nitrogen and stored for later use.

### Tissue Culture

#### Growth of human MCC cell lines

Human MCC cell lines UM-MCC52 (VP-MCC) and UM-MCC50 (VN-MCC) were established in the lab and maintained in an optimized Self Renewal Medium (SRM) as previously described (23).

#### Establishment of mouse MCC cell lines

Four mouse MCC cell lines were established from tumor-bearing SLAP2 mice as previously described for human MCC tumor lines (23). Briefly, a 1-2mm tissue fragment was excised from freshly harvested mouse tumors and finely minced into smaller fragments with scalpel blades. Dissociated tissue was resuspended in optimized SRM supplemented with 4μg/mL doxycycline (Fisher Scientific) to maintain transgene expression.Cultures were monitored by phase-contrast microscopy and supplemented with fresh media as floating cell aggregates expanded. After 1-2 weeks, suspension cultures were moved to clean wells to exclude adherent populations and vigorously triturated to physically separate cells into smaller aggregates. Cell lines were initially passaged 1:2 to maintain a relatively high cell density, or spent media was removed and replaced with fresh medium every 3-5 days as needed. Cells were maintained at 37°C in a humidified, 5% CO2 atmosphere.

Cell culture supernatants were tested for the presence of mycoplasma by the UMich Transgenic Animal Model Core using the PCR-based Lonza MycoAlert Kit, and short tandem repeat (STR) analysis was performed by LabCorp (Burlington, NC) to establish a unique numerical profile for novel cell lines from independent mice.

### Allograft Mouse Models

#### Development of allograft tumors

Mouse MCC cell lines established from SLAP2 tumors (n=4) were injected SQ at 1.0 x 10^6^ cells in 1:1 volume with Matrigel:HBSS in 100μL total volume into the dorsal flanks of ∼8-week-old female immune competent C57BL/6J mice (The Jackson Laboratory Stock No. 000664) to screen for tumorigenicity *in vivo*. Two of the lines that showed 100% penetrance as SQ allografts were additionally injected intradermally (1.0 x 10^6^ cells in 50μL HBSS) into C57BL/6J mice to screen for growth in the dermis, an orthotopic location. Mice were maintained on chow supplemented with doxycycline (200μg/kg, Bioserv Inc.), beginning 3 days prior to cell implantation to maintain transgene expression. Tumor growth kinetics were assessed using calipers to estimate tumor volume (volume=LxW^2^/2) and tumors were measured twice and volumes averaged. Mouse cell line mMCC2 that grew reliably as an orthotopic allograft was used for future studies.

#### Experimental assay for distant metastases

To assess the potential for distant metastases, 1.0 x 10^6^ mMCC2 cells in 100μL HBSS were injected into the tail veins of C57BL/6J female mice. Mice were monitored weekly for visible signs of distress or impaired respiration and harvested at 6 weeks post cell implantation when mice exhibited distended abdomens. Upon dissection, livers, lungs, and lymph nodes were harvested for histological analysis. Livers were weighed prior to fixation.

### Therapeutic treatment strategies of MCC mouse allografts

C57BL/6J mice maintained on doxycycline chow were injected intradermally with 1.0 x 10^6^ mMCC2 cells in 50μL HBSS. At approximately 3-4 weeks post injection, when tumors achieved an average volume of ∼150 mm^3^, mice were randomized into groups and treatments initiated.

#### Immune checkpoint inhibition (ICI)

Treatments with *InVivo*Plus anti-mouse PD-1 (CD279) (Clone J43; BioXCell Cat. No. BP0033-2) were dosed intraperitoneally (IP) twice a week at 200µg/mouse following dilution in saline solution for the duration of treatment experiments.Control mice received the isotype control *InVivo*MAb polyclonal Armenian hamster IgG (BioXCell Cat. No.BE0091) in a similar manner.

#### LSD1-targeted therapy

The LSD1 inhibitor bomedemstat dihydrochoride (MedChemExpress HY-109169C) stock was prepared in DMSO at 250 mg/ml, aliquoted, and stored at -80°C. Working stock was prepared fresh daily via sonication in resuspension buffer (sterile water with 0.5% w/v Tween-80 and 0.5% w/v carboxymethyl-cellulose sodium salt) and dosed daily via oral gavage at 45 mg/kg for 16 days. Control mice were dosed with resuspension buffer only.

#### Combination therapy

Mice were treated as described above but with both anti-PD-1 or bomedemstat. Single treatment groups analyzed in combination studies were also dosed with either resuspension buffer daily via oral gavage or twice a week with control IgG IP. Vehicle-treated groups received both. Mice were treated for 16 consecutive days followed by collection of tumors and lymph nodes for analysis.

### Immunostaining

Tissue in paraffin blocks was sectioned at 5µm, deparaffinized, and rehydrated for immunohistochemical (IHC) or immunofluorescent (IF) staining. Antigen retrieval was carried out in citrate-based buffer (0.01 mol/L citric acid, pH 6.8) or Ready-to-use Citra Plus solution (BioGenex, CA) for 15 minutes at 100^°^ C. Keratin (K) 20 IHC staining required an additional pepsin solution (abcam) antigen retrieval step for 10 minutes at room temperature (1:1 with PBS) following citrate antigen retrieval. For IHC, endogenous peroxidases were first quenched with 3% hydrogen peroxide. Frozen OCT tissue blocks were sectioned and air dried for 15 minutes followed by washing with PBS. Blocking in 5% donkey serum or M.O.M. mouse Ig blocking reagent (Vector Laboratories, Burlingame, CA) for mouse primary antibodies was carried out for 1 hour, followed by incubation of all primary antibodies overnight in a humidified chamber at 4^°^C. Details of primary antibodies including LTAg, ATOH1, K8, K20, Ki67, ISL1, INSM1, SOX2, RCOR2, synaptophysin (SYN), CD3, CD8, and F4/80 are listed in Supplementary Table 1. Bound antibodies were detected with appropriate biotinylated or fluorophore-conjugated (AlexaFluor488, AlexaFlour594, or AlexaFluor555, from Jackson ImmunoResearch Laboratories or Invitrogen) secondary antibodies diluted at 1:300. IHC detection was carried out with the VectaStain ABC or M.O.M. Peroxidase Immunodetection Kits (Vector Laboratories) using SigmaFast diaminobenzidene as a substrate. Hematoxylin was used for nuclear (DNA) counterstaining followed by mounting with Permount (Thermo Fisher Scientific). IF-stained sections were counterstained with DAPI (Sigma-Aldrich) and mounted with ProLong Gold antifade reagent (Invitrogen) for visualization.

### Immunoblotting

Total cell line lysates were obtained by Laemmli extraction, protein was quantified by standard Bradford method using Bio-Rad Protein Assay dye reagent (Bio-Rad Laboratories, CA) and separated on 4-20% gradient SDS-polyacrylamide gels and transferred to Immobilon-P membranes (Millipore, MA). The MCPyV LTAg (CM2B4; Santa Cruz Biotechnology) and ATOH1 (Proteintech) antibodies were used at 1:500 and 1:1000, respectively. Actin (Sigma-Aldrich) was used at 1:10,000 as a loading control. Detection was carried out with SuperSignal West Pico substrate (Thermo Scientific).

### Whole exome sequencing

DNA was isolated from frozen mouse MCC tumors and matched livers of SLAP2 mice using the Qiagen DNeasy Kit and 0.5-1.0 µg of DNA submitted for library preparation and next generation sequencing (NGS) with the NovaSeq S4 (mean on-target coverage depth = 300x) by the BRCF Advanced Genomics Core at the University of Michigan.

#### Tumor mutational burden

Sequencing reads were aligned to the GRCm38 (mm10) primary assembly with BWA-MEM, duplicate-marked (Sentieon Dedup, equivalent to Picard MarkDuplicates), and locally realigned around known Mouse Genomes Project (MGP v6) indels. For each tumor, somatic single-nucleotide variants and short indels were called against its matched germline normal, DNA from the same animal’s liver, using Sentieon TNScope (--min_tumor_allele_frac 0.0075, --min_init_tumor_lod 2.5, --max_fisher_pv_active 0.05, --assemble_mode 4, --trim_soft_clip), with Sentieon DNAscope (--ploidy 4) providing paired germline genotypes and dbSNP/Ensembl mouse variation (build 102) marking known sites. The two lines without a matched liver were instead called against a pooled normal built from all cohort liver samples and were excluded from TMB quantification. Variants were annotated with Ensembl VEP (GRCm38, cache v102). To remove residual germline, strain-background (transgenic vs. C57BL/6), and recurrent-artifact calls, variants were filtered against a cohort panel of normals constructed by TNScope comparison of the pooled tumor-type liver samples, retaining only variants supported by ≤3 reads in the panel; surviving calls were further restricted to TNScope-PASS variants with a moderate- or high-impact coding consequence. TMB was calculated as the number of nonsynonymous coding somatic variants per megabase of callable exonic target territory within the GRCm38 Twist Mouse Exome (rev1) panel.

#### Fraction of genome altered

Copy-number profiles were derived from the whole-exome data using CNVEX. Tumor purity and average ploidy were estimated from the segmented log-ratio and B-allele frequency data, and all tumors were found to be baseline diploid. Segment copy numbers were then converted to integer absolute copy numbers using the estimated purity, and FGA was computed as the proportion of the callable genome whose absolute copy number deviated from the diploid baseline (CN ≠ 2). FGA was therefore based on purity-corrected absolute copy-number states rather than a fixed log-ratio threshold, which avoids purity-dependent miscalling of altered segments.

Human data used to compare TMB in VN-MCC and VP-MCC to mouse MCC tumors was previously generated and described (5).

### Serologic assay

Serum was isolated following cardiac puncture and terminal exsanguination of tumor-bearing MCC mice using BD microtainer gel tubes for serum collection as per manufacturer’s instructions. A quantitative multiplex binding assay for detection of serum antibodies against MCPyV TAgs was carried out at the Fred Hutchinson Cancer Center. MCPyV T antigen fusion proteins, in which glutathione S-transferase (GST) was added to the N-terminus and an 11 amino acid epitope tag added to the C-terminus were used for all antibody binding assays. The T antigen constructs included LTAg and sTAg, and the region of LT and sT antigens which are shared, common T (CT), as previously described (56). Serial dilutions of sera beginning at 1:50 were tested. The subtracted median fluorescent intensity (MFI) values for LT and sT antigens were used to generate titration curves on GraphPad Prism (GraphPad Prism Software Inc, La Jolla, CA) using the Sigmoidal dose response (with variable slope) program. Pre-incubation with either the GST-CT fusion protein or with a monoclonal antibody that recognized the CT region, CM2B4 (Sana Cruz Biotechnology) nearly completely inhibited the reactivity to LT and sT. Sera from *K5-SLAP2* mouse littermates genotyped negative for *tetO-sT/tLT* transgenes were used as controls for sera collection.

### Statistics

Statistical analysis for tumor growth kinetics, including mean tumor volumes and error bars (SEM) were calculated in Excel for Microsoft 365 MSO (Version 2603). Mean tumor volumes and p values (two-tailed unpaired t test using the Holm-Sidak method) were calculated in GraphPad Prism (v10.4.1). Significance was designated as follows: *P < 0.05*, **P < 0.01, ***P< 0.001, and ****P < 0.0001.

To determine differences in longitudinal tumor growth rate, a linear mixed model (32, 33) was used to estimate the fitted line for each treatment group from individual tumor responses (**Fig. 5b** solid and dashed lines, respectively). The model had an outcome of log tumor volume and included fixed effects for time (in days), treatment group, and a time-group interaction, and a random intercept and slope for each ID. Analyses were performed in R (version 4.3.2) using the lmerTest package.

## Acknowledgements

We thank personnel in the Dlugosz and Wong labs for helpful comments throughout the course of these studies. This work was supported in part by the National Cancer Institute (P30 CA046592) (MEV, PWH, MC, AF, AAD), (R01 CA189352, R01 CA241947) (MEV, AAD),(P01 CA225517) (MEV, DAG, AAD); National Institute of Arthritis and Musculoskeletal and Skin Diseases (P30 AR075043) (MEV, LJS, AAD); the Linkou Chang Gung Memorial Hospital Research Scholar Program (PWH); the Autenrieth Merkel Cell Fund (AAD); and A. Alfred Taubman Medical Research Institute, Michigan Medicine, Taubman Institute Innovation Program (MEV, LJS, PWH, AF, MC, AAD).

