## Supplementary Data for "An immunocompetent Merkel cell carcinoma model for preclinical studies"

### Slide 1
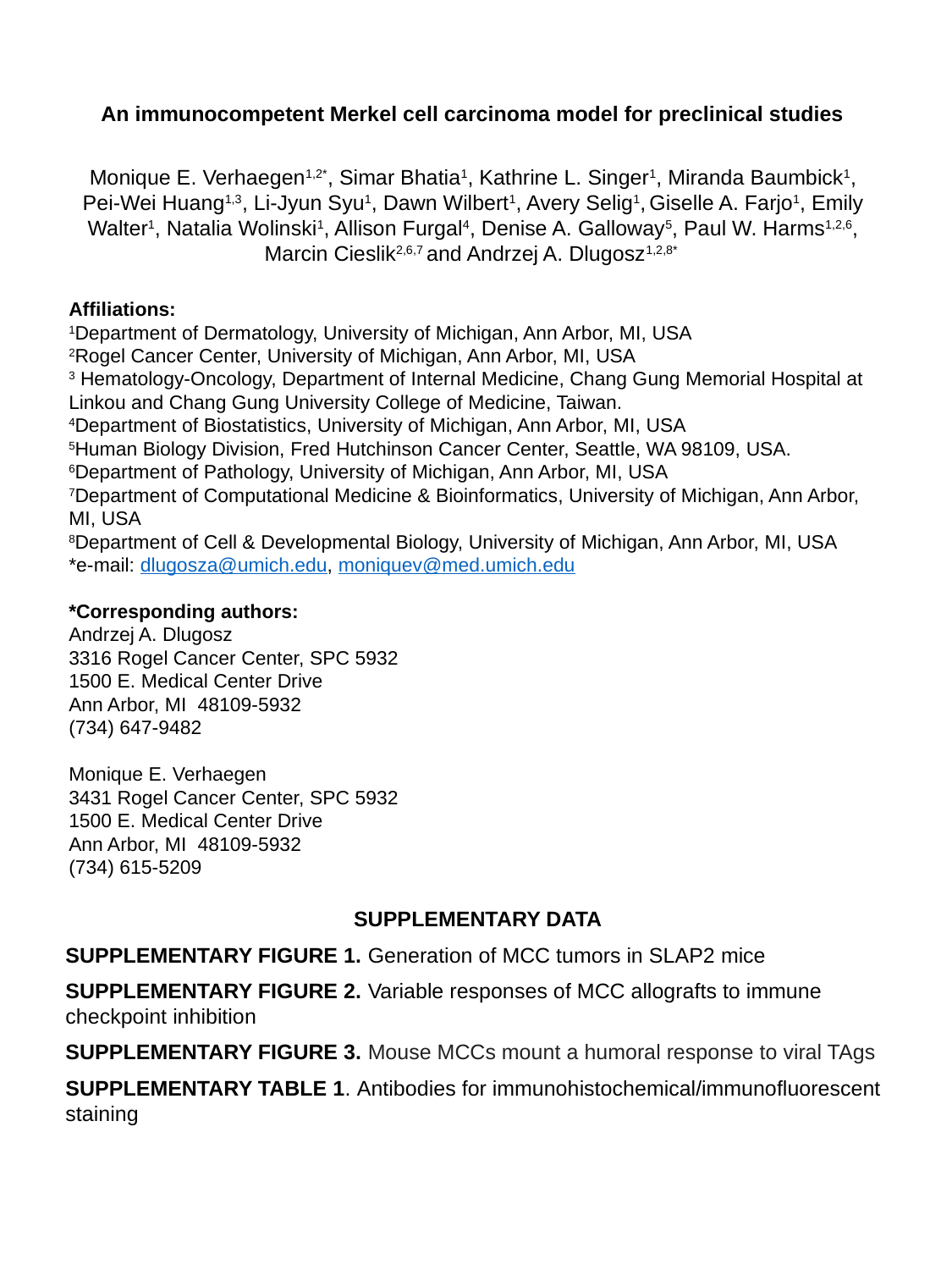

An immunocompetent Merkel cell carcinoma model for preclinical studies
Monique E. Verhaegen^1,2*^, Simar Bhatia^1^, Kathrine L. Singer^1^, Miranda Baumbick^1^, Pei-Wei Huang^1,3^, Li-Jyun Syu^1^, Dawn Wilbert^1^, Avery Selig^1^, Giselle A. Farjo^1^, Emily Walter^1^, Natalia Wolinski^1^, Allison Furgal^4^, Denise A. Galloway^5^, Paul W. Harms^1,2,6^, Marcin Cieslik^2,6,7^ and Andrzej A. Dlugosz^1,2,8*^
Affiliations:
^1^Department of Dermatology, University of Michigan, Ann Arbor, MI, USA
^2^Rogel Cancer Center, University of Michigan, Ann Arbor, MI, USA
^3^ Hematology-Oncology, Department of Internal Medicine, Chang Gung Memorial Hospital at Linkou and Chang Gung University College of Medicine, Taiwan.
^4^Department of Biostatistics, University of Michigan, Ann Arbor, MI, USA
^5^Human Biology Division, Fred Hutchinson Cancer Center, Seattle, WA 98109, USA.
^6^Department of Pathology, University of Michigan, Ann Arbor, MI, USA
^7^Department of Computational Medicine & Bioinformatics, University of Michigan, Ann Arbor, MI, USA
^8^Department of Cell & Developmental Biology, University of Michigan, Ann Arbor, MI, USA
*
*Corresponding authors:
Andrzej A. Dlugosz
3316 Rogel Cancer Center, SPC 5932
1500 E. Medical Center Drive
Ann Arbor, MI 48109-5932
(734) 647-9482
Monique E. Verhaegen
3431 Rogel Cancer Center, SPC 5932
1500 E. Medical Center Drive
Ann Arbor, MI 48109-5932
(734) 615-5209
SUPPLEMENTARY DATA
SUPPLEMENTARY FIGURE 1. Generation of MCC tumors in SLAP2 mice
SUPPLEMENTARY FIGURE 2. Variable responses of MCC allografts to immune checkpoint inhibition
SUPPLEMENTARY FIGURE 3. Mouse MCCs mount a humoral response to viral TAgs
SUPPLEMENTARY TABLE 1. Antibodies for immunohistochemical/immunofluorescent staining

### Slide 2
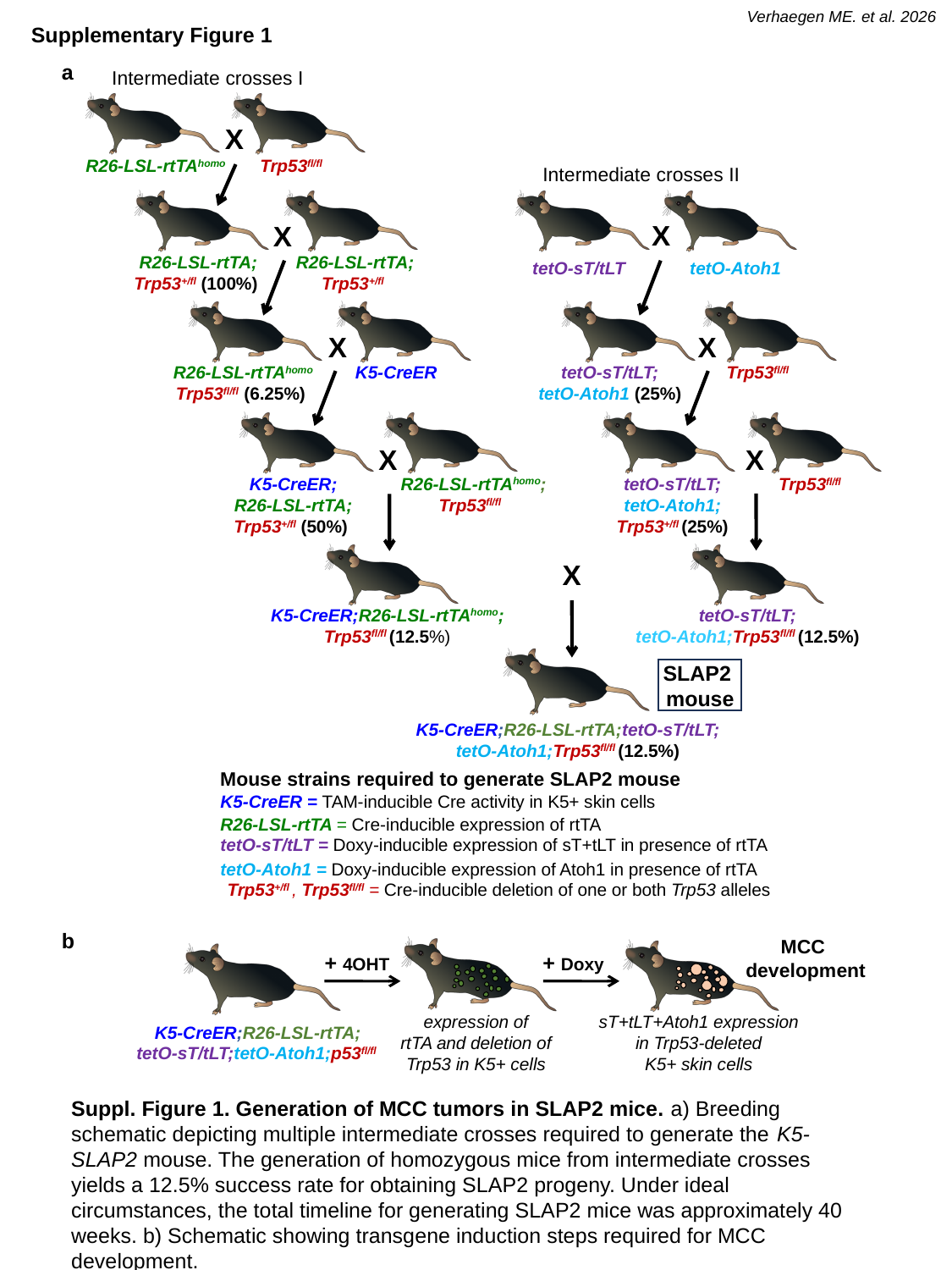

Verhaegen ME. et al. 2026
Supplementary Figure 1
a
Intermediate crosses I
X
R26-LSL-rtTA^homo^
Trp53^fl/fl^
Intermediate crosses II
X
tetO-sT/tLT
tetO-Atoh1
X
tetO-sT/tLT;
tetO-Atoh1 (25%)
Trp53^fl/fl^
X
tetO-sT/tLT;
tetO-Atoh1;
Trp53^+/fl^ (25%)
Trp53^fl/fl^
tetO-sT/tLT;
tetO-Atoh1;Trp53^fl/fl^ (12.5%)
X
R26-LSL-rtTA;
Trp53^+/fl^ (100%)
R26-LSL-rtTA;
Trp53^+/fl^
X
R26-LSL-rtTA^homo^
Trp53^fl/fl^ (6.25%)
K5-CreER
X
K5-CreER;
R26-LSL-rtTA;
Trp53^+/fl^ (50%)
R26-LSL-rtTA^homo^;
Trp53^fl/fl^
X
K5-CreER;R26-LSL-rtTA^homo^;
Trp53^fl/fl^ (12.5%)
SLAP2
mouse
K5-CreER;R26-LSL-rtTA;tetO-sT/tLT;
tetO-Atoh1;Trp53^fl/fl^ (12.5%)
Mouse strains required to generate SLAP2 mouse
K5-CreER = TAM-inducible Cre activity in K5+ skin cells
R26-LSL-rtTA = Cre-inducible expression of rtTA
tetO-sT/tLT = Doxy-inducible expression of sT+tLT in presence of rtTA
tetO-Atoh1 = Doxy-inducible expression of Atoh1 in presence of rtTA
Trp53^+/fl^ , Trp53^fl/fl^ = Cre-inducible deletion of one or both Trp53 alleles
b
MCC
development
+ 4OHT
+ Doxy
expression of
rtTA and deletion of Trp53 in K5+ cells
sT+tLT+Atoh1 expression
in Trp53-deleted
K5+ skin cells
K5-CreER;R26-LSL-rtTA;
tetO-sT/tLT;tetO-Atoh1;p53^fl/fl^
Suppl. Figure 1. Generation of MCC tumors in SLAP2 mice. a) Breeding schematic depicting multiple intermediate crosses required to generate the K5-SLAP2 mouse. The generation of homozygous mice from intermediate crosses yields a 12.5% success rate for obtaining SLAP2 progeny. Under ideal circumstances, the total timeline for generating SLAP2 mice was approximately 40 weeks. b) Schematic showing transgene induction steps required for MCC development.

### Slide 3
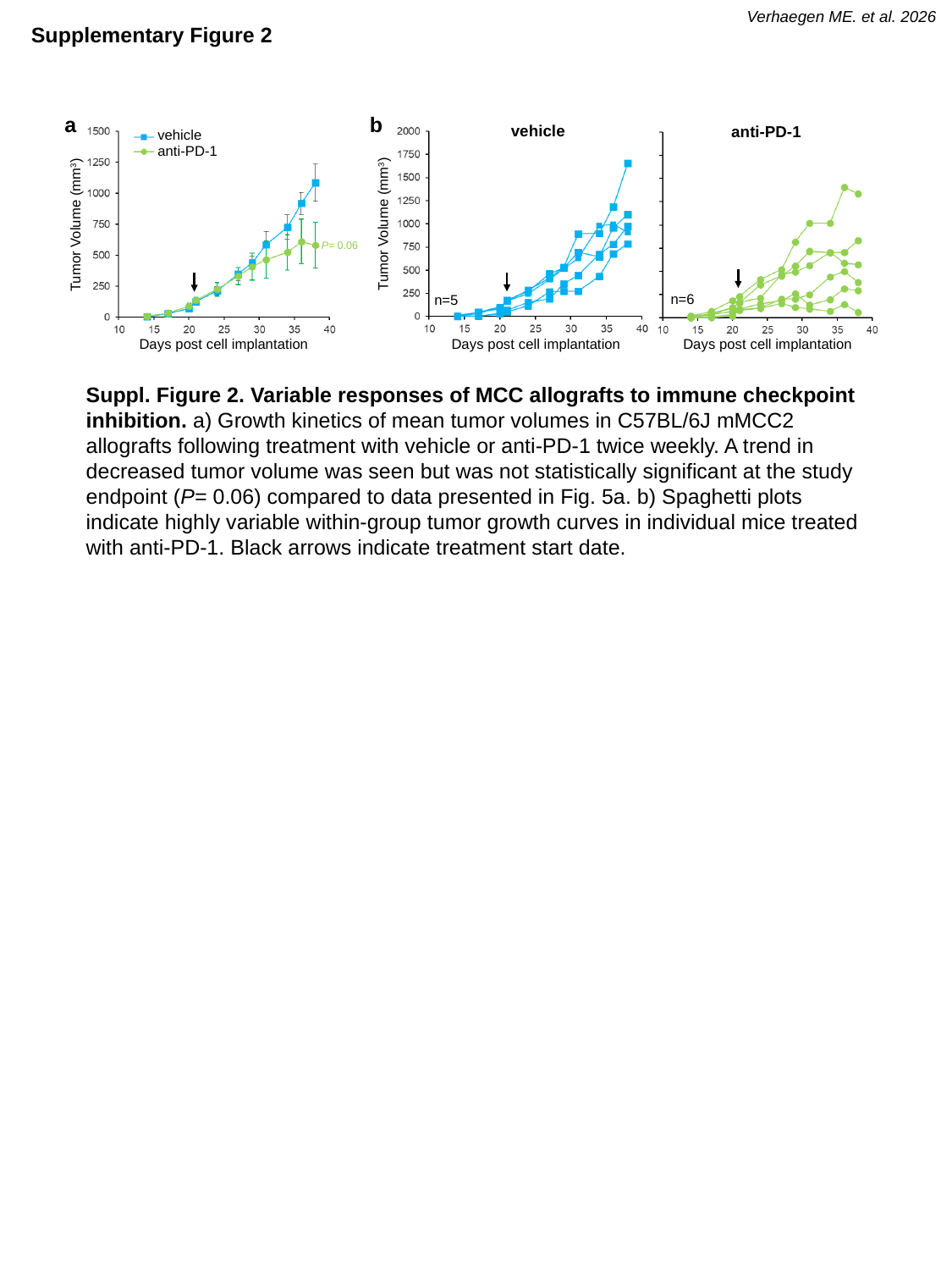

Verhaegen ME. et al. 2026
Supplementary Figure 2
a
b
vehicle
anti-PD-1
vehicle
anti-PD-1
Tumor Volume (mm^3^)
Tumor Volume (mm^3^)
P= 0.06
n=6
n=5
Days post cell implantation
Days post cell implantation
Days post cell implantation
Suppl. Figure 2. Variable responses of MCC allografts to immune checkpoint inhibition. a) Growth kinetics of mean tumor volumes in C57BL/6J mMCC2 allografts following treatment with vehicle or anti-PD-1 twice weekly. A trend in decreased tumor volume was seen but was not statistically significant at the study endpoint (P= 0.06) compared to data presented in Fig. 5a. b) Spaghetti plots indicate highly variable within-group tumor growth curves in individual mice treated with anti-PD-1. Black arrows indicate treatment start date.

### Slide 4
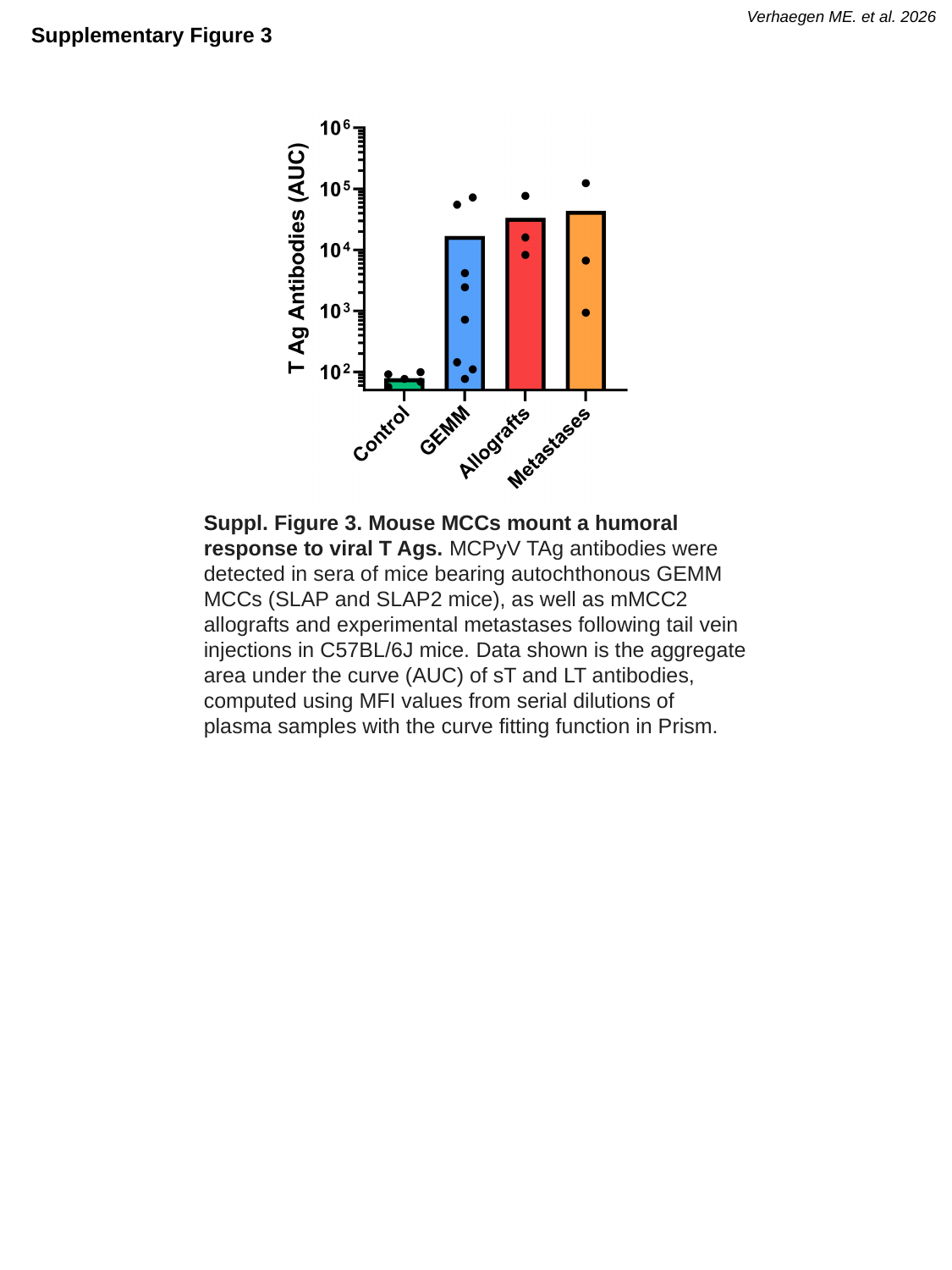

Verhaegen ME. et al. 2026
Supplementary Figure 3
Suppl. Figure 3. Mouse MCCs mount a humoral response to viral T Ags. MCPyV TAg antibodies were detected in sera of mice bearing autochthonous GEMM MCCs (SLAP and SLAP2 mice), as well as mMCC2 allografts and experimental metastases following tail vein injections in C57BL/6J mice. Data shown is the aggregate area under the curve (AUC) of sT and LT antibodies, computed using MFI values from serial dilutions of plasma samples with the curve fitting function in Prism.

### Slide 5
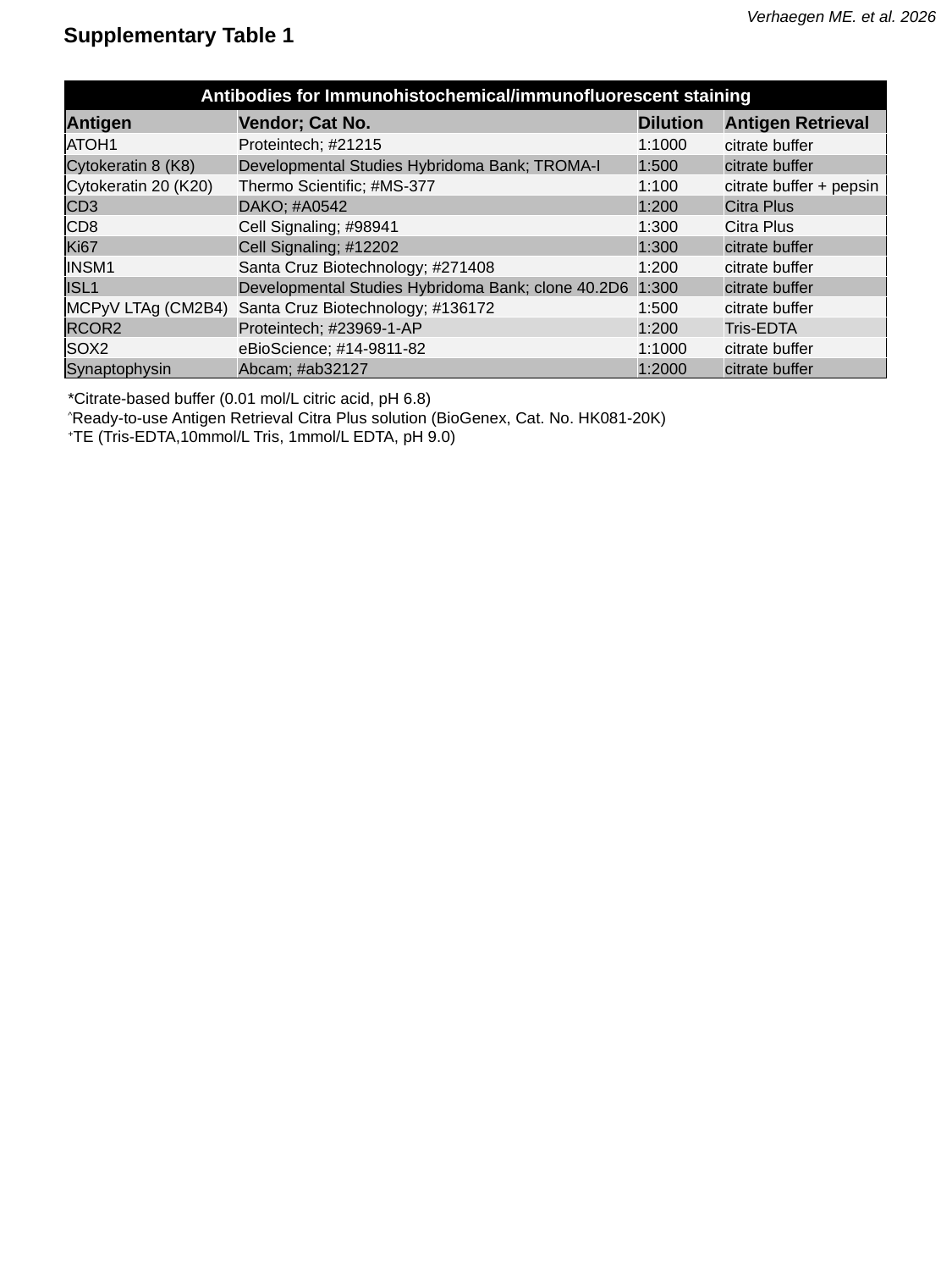

Verhaegen ME. et al. 2026
Supplementary Table 1
| Antibodies for Immunohistochemical/immunofluorescent staining | | | |
| --- | --- | --- | --- |
| Antigen | Vendor; Cat No. | Dilution | Antigen Retrieval |
| ATOH1 | Proteintech; #21215 | 1:1000 | citrate buffer |
| Cytokeratin 8 (K8) | Developmental Studies Hybridoma Bank; TROMA-I | 1:500 | citrate buffer |
| Cytokeratin 20 (K20) | Thermo Scientific; #MS-377 | 1:100 | citrate buffer + pepsin |
| CD3 | DAKO; #A0542 | 1:200 | Citra Plus |
| CD8 | Cell Signaling; #98941 | 1:300 | Citra Plus |
| Ki67 | Cell Signaling; #12202 | 1:300 | citrate buffer |
| INSM1 | Santa Cruz Biotechnology; #271408 | 1:200 | citrate buffer |
| ISL1 | Developmental Studies Hybridoma Bank; clone 40.2D6 | 1:300 | citrate buffer |
| MCPyV LTAg (CM2B4) | Santa Cruz Biotechnology; #136172 | 1:500 | citrate buffer |
| RCOR2 | Proteintech; #23969-1-AP | 1:200 | Tris-EDTA |
| SOX2 | eBioScience; #14-9811-82 | 1:1000 | citrate buffer |
| Synaptophysin | Abcam; #ab32127 | 1:2000 | citrate buffer |
*Citrate-based buffer (0.01 mol/L citric acid, pH 6.8)
^^^Ready-to-use Antigen Retrieval Citra Plus solution (BioGenex, Cat. No. HK081-20K)
^+^TE (Tris-EDTA,10mmol/L Tris, 1mmol/L EDTA, pH 9.0)
